# mRNAid, an Open-Source Platform for Therapeutic mRNA Design and Optimization Strategies

**DOI:** 10.1101/2022.04.04.486952

**Authors:** Nikita Vostrosablin, Shuhui Lim, Pooja Gopal, Kveta Brazdilova, Sushmita Parajuli, Xiaona Wei, Anna Gromek, Martin Spale, Anja Muzdalo, Constance Yeo, Joanna Wardyn, Petr Mejzlik, Brian Henry, Anthony W. Partridge, Danny A. Bitton

**Author notes:** **Corresponding Author: Danny A. Bitton**, Na Valentince 4, FIVE Building, Prague 5 - Smichov, Prague, 150 00, Czech Republic. Equal Contribution (Nikita Vostrosablin, Shuhui Lim, Pooja Gopal).

## Abstract

Recent COVID-19 vaccines unleashed the potential of mRNA-based therapeutics. mRNA optimization is indispensable for reducing immunogenicity, ensuring stability, and maximizing protein output. We present mRNAid, an experimentally validated software for mRNA optimization and visualization that generates mRNA sequences with comparable if not superior characteristics to commercially optimized sequences. To encompass all aspects of mRNA design, we also interrogated the impact of uridine content, nucleoside analogs and UTRs on expression and immunogenicity.

## Main

Synthetic mRNA-based therapeutics continue to revolutionize vaccine development^1^, immunotherapy^2^ and targeted degradation methodologies^3^, advancing the battle of modern medicine against infectious diseases^4^, genetic disorders^5^ and cancer^6^. Irrespective of the various indications and the diverse mechanisms of action of mRNA-based drugs, all derived therapeutics share the same underlying principles. First, the mRNA sequence is designed and optimized *in-silico*. Then, the optimized sequence is transcribed with selected chemical modifications *in-vitro*. Finally, the synthetic transcript is packaged and delivered to the cytoplasm of host cells, where it is translated into a protein that exerts the desired cellular effect. Therefore, *in-silico* mRNA design is undeniably instrumental to the success of any mRNA-based therapeutic. Transcript design is typically initiated with a decoration of the coding sequence (CDS) with flanking 5’ and 3’ UTRs (untranslated regions) and other signals (e.g. translational ramps, miRNA binding sites, etc.) that can improve stability, translation efficiency and enable tissue-specific expression^7,8^. Then, rigorous sequence engineering is required to eliminate immunogenic properties^9^ and further enhance transcript stability and translation^10,11^. During *in-vitro* transcription, chemical modifications such as 5’-cap and nucleoside analogs are incorporated to protect against degradation and evade host immune surveillance^12^. At present, mRNA design is largely dependent on expert knowledge, manual sequence editing, distributed optimization and visualization tools that are often proprietary. Thus, there is no freely available tool specifically tailored for therapeutic mRNA design that combines multiple optimization strategies. Here, we present mRNAid, an open-source, integrated software that bundles several modified and extended algorithms and tools for constraint propagation, sequence optimization and secondary structure visualization. Via an intuitive and user-friendly interface, mRNAid orchestrates simultaneous optimization of several sequence and structural properties including codon usage, GC content, minimum free energy (MFE), uridine depletion and exclusion of specific motifs and/or rare codons, thereby providing a powerful platform for therapeutic mRNA design.

The DNA Chisel framework was chosen as the core backbone for sequence optimization in mRNAid^13^ (Supplementary Figure S1). DNA Chisel permits global and local optimization of hard and soft constraints, which in turn enables the adjustment of desired sequence properties along the entire transcript. Hard constraints refer to criteria that must be satisfied in the final sequence, whereas soft constraints refer to criteria whose score must be maximized. Furthermore, the ability to flexibly define new constraints makes DNA Chisel an ideal sandbox for probing the effect of a multitude of sequence properties on stability and expression.

mRNAid piggybacks on several hard constraints implemented in DNA Chisel that enforce global GC content and translation and avoid rare codons and specific motifs. mRNAid extends this list with an important uridine depletion constraint that avoids codons with uridine at their third position, which reportedly improves expression and reduces immunogenicity^9^.

Codon usage optimization aims to improve expression by systematic replacement of synonymous codons based on the organism’s codon frequency table. Numerous proprietary and freely available codon optimization algorithms have been reported to date^10^ and many show a strong preference towards the Codon Adaptation Index (CAI)^14^ since it highly correlates with gene expression. DNA Chisel includes CAI and Matched Codon Usage^15^ optimizations as soft constraints. We also implemented the dinucleotide and codon-pair usage optimizations based on the CoCoPUTs database that have been shown to affect translation fidelity and efficiency^16^. There is a significant codon-pair usage bias in all three domains of life and between lowly and highly expressed proteins within a species^17^ that cannot be explained by individual codon bias, pointing towards a distinct mechanism of translation modulation. It has been suggested that codon-pair effects on translation rate may be mediated by interactions of adjacent aminoacyl-tRNA molecules bound to ribosomes^18^. To account for structural properties, we also incorporated the Vienna-RNA MFE optimization^19^ and the correlated stem–loop prediction approach^20^, given the pivotal role that mRNA secondary structures play in regulating translation efficiency. Multiple studies reported correlation between highly structured features in the coding sequence (CDS) and functional mRNA half-life^21, 22^. In contrast, the region around the translation start site is less structured in highly expressed genes^23^, which presumably facilitates ribosome loading and prevents jamming^24^. Thus, different transcript regions possess different structural properties that must be reflected in the optimization strategy. Furthermore, transcripts are often fused to already optimized UTRs, obviating the need for optimization of these regions. Given the above, and the fact that MFE optimization is extremely computationally expensive, we provide users with the flexibility to optimize MFE within an adjustable window in the 5’-end of the CDS, while accounting for global MFE in our ranking approach. To this end, we ensure that the combination of multiple constraints generates optimal sequence properties by applying a novel scoring formula that ranks sequences based on weighted scores of uridine depletion, GC content, CAI, local 5’-end and global MFEs.

Next, we selected five distinct optimization strategies in mRNAid that are based on a codon optimization approach coupled with uridine depletion, GC and MFE optimizations (Strategies 1-5: dinucleotide, matched codon-pair usage, matched codon usage with or without uridine depletion and CAI, respectively). To experimentally validate these strategies, we adopted NanoLuciferase-PEST (NLuc-PEST) as the reporter system since the short half-lives of the individually produced luciferase proteins prevent confounding effects on interrogation of mRNA properties as they relate to translation efficiency or mRNA stability. Indeed, these effects can be conveniently measured through kinetic tracking of the target protein’s luminescence. Area under the curve (AUC) and luminescence at 48 hours (RLU @ 48h) are represented as indicators of total protein output and functional mRNA stability respectively, given that protein sequence and other mRNA features are kept constant. A de-optimized mRNA version of NLuc-PEST encoded by the least frequent codon for each amino acid (Rare, red, Figure 1A-C) was used as the input for mRNAid and the top 6 ranked mRNA sequences generated under each software setting were selected (Supplementary Table S1 and S2). These were benchmarked against a proprietary sequence from Promega (Promega, blue, Figure 1A-C) as the codon-optimized control.

**Figure 1.**
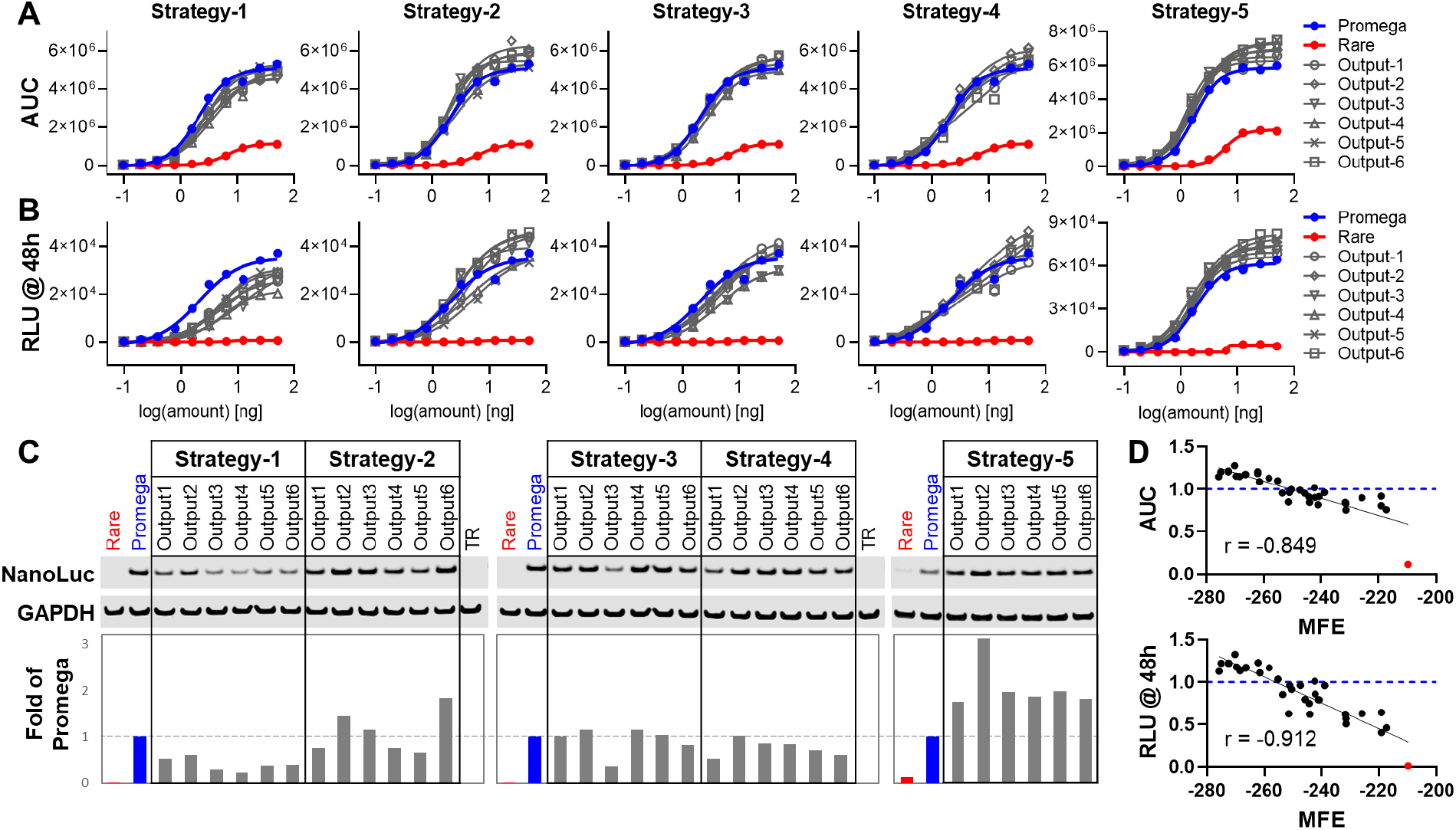
Impact of various sequence optimization strategies in mRNAid on NanoLuc-PEST expression. Effect of 5 different sequence optimization strategies on NanoLuc-PEST expression in MIA PaCa-2 was represented as (A) area under the curve, AUC of luminescence over 48 hr and (B) relative luminescence, RLU at 48 hr post-transfection. Rare, red denotes the de-optimized input and Promega, blue denotes the codon-optimized control. The top 6 outputs from each strategy were tested. (C) Western blot analysis of the sequence variants from above 24 hr post-transfection in MIA PaCa-2 cells. GAPDH was used as a loading control. Band intensities for NanoLuc were normalized to GAPDH and represented as fold change over the Promega control. (D) Correlation plots for AUC (top) and RLU @ 48 hr (bottom) versus MFE (kcal/mol). AUC and RLU @ 48 hr were determined for the 6.25 ng mRNA dose in (A and B) and represented as fold change over the Promega control (blue dashed line). The red dot denotes the de-optimized Rare input. Pearson r is indicated as determined by GraphPad Prism.

All 30 mRNAid-optimized sequences resulted in significantly higher expression relative to the de-optimized input (Figure 1A-C), attesting to the robustness of the tool. Encouragingly, out of these 30 sequences, 8 (30%) surpassed and 14 (47%) matched the expression of the codon-optimized Promega control (Figure 1C). Among the five optimization strategies, all 6 sequences that were optimized by Strategy-5 (CAI optimization) expressed better than Promega and exhibited the highest GC content, lowest global MFE and lowest uridine content (Figure 1A-C, Supplementary Table S2). Similarly, sequences that were optimized by Strategy-2 (matched codon-pair optimization) exhibited comparable (4/6) or better expression (2/6) than Promega. The importance of codon-pair context for enhanced gene expression was also recognized in previous studies^25^. Conversely, all the 6 sequences that were optimized by Strategy-1 (dinucleotide optimization) expressed poorer than Promega (Figure 1A-C, Supplementary Table S2). An *in-silico* optimization of the native SARS-CoV-2 surface glycoprotein gene using mRNAid revealed that among all optimization approaches Strategy-5 (CAI optimization) has generated transcripts with the highest similarity to the assembled Moderna and Pfizer/BioNtech vaccine sequences (∼96.5% and ∼90.8% similarity, respectively, Supplementary Table S1)^26, 27^, suggesting that a similar optimization strategy was employed for their design. Further analyses revealed that both AUC and RLU @ 48hr were strongly correlated with global MFE (r = -0.85 and r = -0.91, respectively, Figure 1D), in line with previous reports on the impact of secondary structures on mRNA half-lives and ultimately final protein output^21^. Correlations were also observed with GC (%) and U-ratio, but not 5’-MFE (Supplementary Figure S2). These findings highlight the importance of combining a multitude of hard, soft, local and global constraints to achieve balanced sequence and structural properties that cooperatively define total protein expression.

We next explored additional features that can be incorporated into mRNAid-optimized sequences to yield optimal therapeutic mRNAs. The discovery that uridine analogs dramatically reduce immune stimulation^9^ and increase protein production from synthetic mRNA^12^ marked a breakthrough in mRNA-based therapeutics. We evaluated the impact of substituting uridine (U) with pseudouridine (pU), 5-methoxyuridine (5moU) or N1-methylpseudouridine (N1m) on NLuc-PEST protein expression and pro-inflammatory cytokine (IFN-β) release in human BJ fibroblasts. For the de-optimized input (Rare) and codon-optimized control (Promega) sequences, pU and N1m modifications significantly improved protein expression as compared to U (Figure 2A), in line with previous reports. Instead of improving expression, U substitution with 5moU reduced protein output from the Promega sequence. Out of 30 mRNAid sequences, we picked the best expressed sequence (Strategy-5:output-2, Figure 1C) and were able to recapitulate this improvement compared to Rare and Promega in BJ fibroblast using U (black, Figure 2A). However, protein levels were not further enhanced using uridine analogs in Strategy-5:output-2 sequence. We postulate that since Strategy-5:output-2 sequence had the lowest uridine content (14% in Strategy-5:output-2; 21% in Promega; 27% in Rare), the effect of uridine substitution on expression may be minimal. In terms of immunogenicity, unmodified mRNAs (U) caused the highest cytokine release as expected (comparable to positive control poly I:C), followed by pU, N1m and 5moU (Figure 2B). Among the 3 sequences tested, mRNAid-optimized Strategy-5:output-2 was consistently less immunogenic than Rare and Promega (Figure 2B) under conditions where IFN-β was within detectable range. Thus, our findings demonstrate how sequence optimization combined with incorporation of uridine analogs can reduce undesired innate immune responses while maintaining high target protein expression.

**Figure 2.**
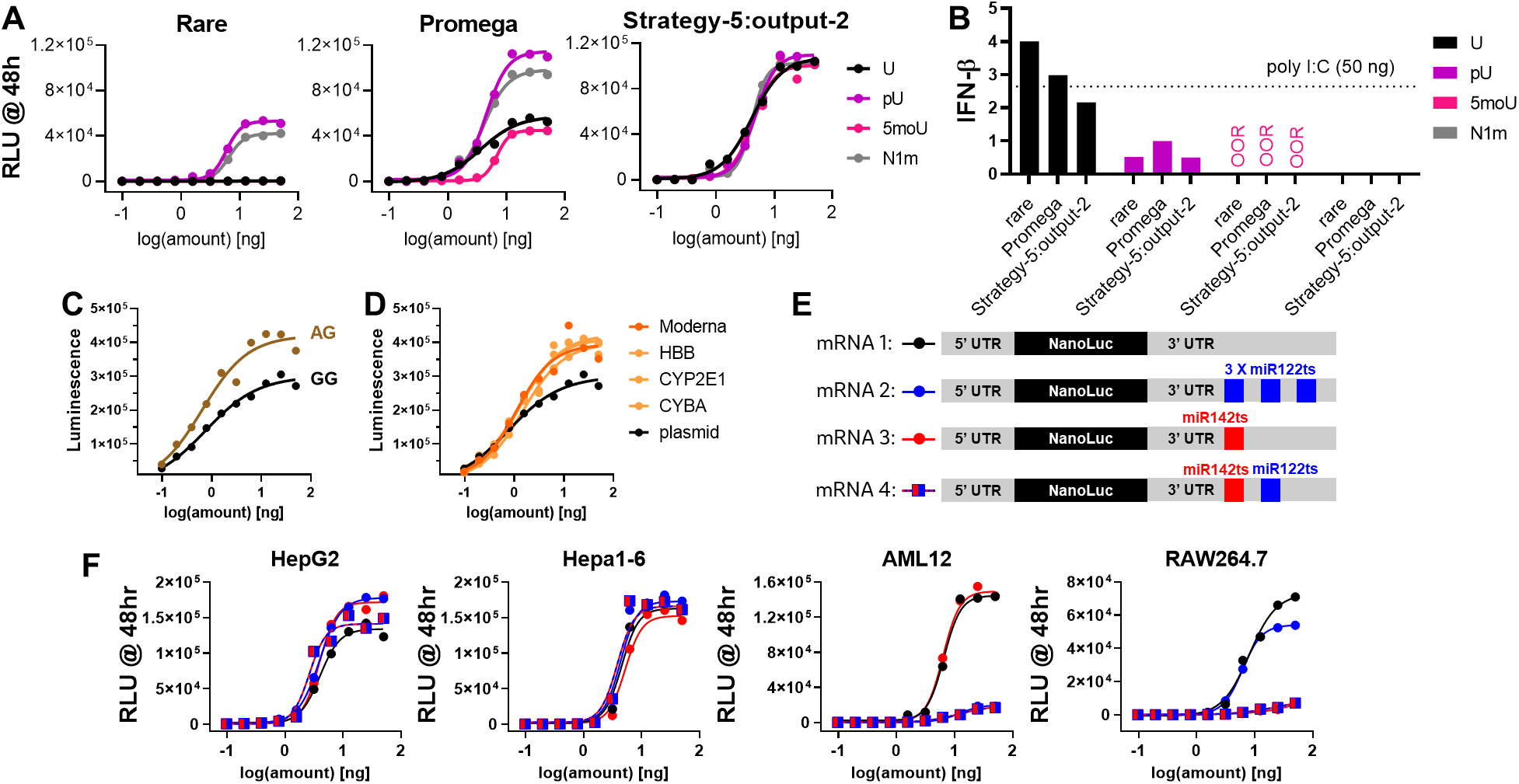
Additional ways to engineer mRNA for therapeutic use. (A) Effect of modified nucleotides on NanoLuc-PEST expression in BJ fibroblasts. U, uridine; pU, pseudouridine; 5moU, 5-methoxyuridine; N1m, N1-methyl-pseudouridine. (B) Effect of modified nucleotides on innate immune activation in BJ fibroblasts. Cytokine release assay 48 hr post-transfection with 50 ng of the indicated mRNAs. IFN-β levels were normalized to the Promega sequence with pU incorporated. OOR, out-of-range. The dashed line represents positive control poly I:C. (C) Effect of AG versus GG initiator sequence after the T7 promoter on NanoLuc protein expression in MIA PaCa-2. (D) Effect of UTRs on NanoLuc protein expression in MIA PaCa-2. (E) Design of NanoLuc mRNAs with miRNA target sites in 3’ UTR. (F) Effect of self-destruct signals on NanoLuc protein expression in the indicated cell type.

mRNAs are capped at the 5’-end to protect against degradation, facilitate ribosome loading and evade innate immune responses^28^. Compared to legacy cap analogs such as ARCA, the proprietary co-transcriptional capping reagent CleanCap AG from TriLink was shown to have higher capping efficiency, increased RNA yield and reduced immunogenicity. We modified the conventional GG initiator sequence after the T7 promoter to AG and synthesized mRNAs using CleanCap AG. Indeed, the AG initiator significantly improved NanoLuc-PEST expression (Figure 2C) and total RNA yield (data not shown). This highlights how protein output can be further boosted and emphasizes on the importance of tailoring template design to the desired capping technique. As noted, the role of 5’ and 3’-UTRs in modulating mRNA stability and translation is well established^22^. To this end, we selected 4 pairs of UTR sequences that have been reported to boost protein expression in human cells (Supplementary Table S3). Indeed, all 4 UTRs increased NanoLuc expression compared to plasmid which are default sequences flanking the CDS (Figure 2D). Together, we present various opportunities to enhance mRNA potency by optimizing key mRNA components.

A therapeutic mRNA should ideally be delivered to and expressed at the target site to minimize toxicities in unintended recipient cells. Jain *et al*. previously demonstrated that microRNAs (miRNA) with disease or tissue-specific expression profiles can be recruited to achieve selective degradation of synthetic mRNAs^7^. For example, miRNA-122 is abundant in normal liver cells but significantly down-regulated in liver carcinoma^29^, while miRNA-142 is predominantly expressed in hemopoietic cells^30^. Using the same strategy, we incorporated miRNA-122 and miRNA-142 target sites to the 3’-UTRs of NanoLuc mRNA to act as self-destruct signals (Figure 2E). Our results confirmed selective mRNA silencing in normal liver (AML12) and blood (RAW264.7) cells by miR122ts and miR142ts respectively, as opposed to unaltered expression in mouse (Hepa1-6) and human (HepG2) liver cancer cell lines (Figure 2F). We further demonstrate that a combination of miR122ts and miR142ts suppressed expression in both AML12 and RAW264.7, highlighting the potential to combine self-destruct signals and possibly other mRNA regulatory elements to refine target protein expression.

In summary, this study represents a first attempt to create a comprehensive playbook for rational design of therapeutic mRNA transcripts. mRNAid is an open-source software that offers advanced sequence and structural optimization strategies that generate transcripts with desired expression properties. We also experimentally demonstrate that incorporation of certain uridine analogs, and inclusion of key mRNA components can further enhance stability, boost protein output, enable tissue-specific expression and mitigate undesired immunogenicity effects.

Despite the encouraging results of mRNAid, it is important to note its limitations. mRNAid does not optimize for MFE along the entire transcript and the thermodynamic parameters of uridine analogs are not accounted for. Yet the experimental data we presented here clearly indicate that global MFE optimization is worthwhile, albeit computationally expensive. However, the flexible backbone of mRNAid offers the greater scientific community the opportunity to make such improvements by adding more efficient optimization strategies as they emerge or by combining existing optimization methods, constraints or other sequence features that may govern stability, immunogenicity, and expression.

## Methods

### Tool architecture

The application consists of several parts, which are containerized and can be easily built and run with ‘docker-compose’ utility (Supplementary Figure S1). The frontend is served as static files with the help of Nginx server. It can also be configured as a reverse proxy. Frontend container communicates with backend through uwsgi protocol. The backend presents a Python Flask application served by uWSGI server. The optimization tasks are handled by celery task queue implemented with Redis in-memory database working as message broker. The mounted volume is used to keep logs of the backend execution. All individual parts reside within separate containers that communicate with each other inside a docker network as represented in Supplementary Figure S1. The user interface is written in React.js and consists of an input form and results page. Input form allows user to select different optimization strategies, set optimization parameters and submit the optimization job. The output form includes visualization of the optimized sequences generated via Forna JavaScript visualization container^19^ combined with MFE mountain-plot and summary of optimized sequence properties. User can export the results in a pdf or an Excel format.

### Optimization strategies

The core of the tool is the freely available sequence optimization framework for Python, DNA Chisel. DNA Chisel allows to use built-in specifications to approach some of the common optimization tasks (like matching target codon usage in host or ensuring correct translation to protein by using only synonymous codons during the optimization, etc.) and it is very flexible with respect to defining completely new optimization specifications. These specifications can either be hard constraints, which cannot be violated in the final sequence, or they can be considered as soft constraints or objectives, whose score is maximized in the final sequence. Some specifications can be used as both constraint and objective depending on user requirements. When multiple objectives are defined in the optimization problem, the total weighted score is maximized.

DNA Chisel solves the constraint satisfaction problem using a combination of constraint propagation and local search methods. The optimization algorithm consists of two main steps, the resolution of all hard constraints and maximization of objectives’ scores with respect to the constraints. The solver reduces the optimization problem to the set of local optimization problems, which are resolved individually. The optimization is performed either by random mutations on the sequence or by exhaustive search through the precomputed mutation space, depending on the size of the latter. During the mRNAid optimization, the tool combines all the specified constraints and objectives together into the optimization problem and calls the DNA Chisel optimization method on it. This procedure is re-executed in parallel until either a user-specified number of sequences are produced or until the number of attempts is exceeded. After the optimization is completed, mRNAid ranks them based on the scoring function, as described below.

In our optimization approach we used following built-in specifications. As hard constraints we used AvoidPattern to ensure that certain motifs are excluded, EnforceGCContent to keep GC levels in a certain boundary across the sequence, AvoidRareCodons to not use rare codons with codon frequency below the threshold and EnforceTranslation to ensure that the optimized sequence is translated back to the same protein as the input. As objectives we used built-in MatchTargetCodonUsage to be as close as possible to the codon usage frequencies in the host and EnforceGCContent to optimize GC content in the sliding window across the sequence. The details of these specifications can be found in the DNAChisel documentation^13^.

We also implemented our own specifications and integrated them into the tool. As hard constraints we implemented UridineDepletion which makes sure that no codons with uridine in the third position are present. We also implemented three new objectives: MatchTargetPairUsage to account for dinucleotide usage, MatchTargetCodonPairUsage to optimize for codon-pair usage frequencies and MinimizeMFE to use different algorithms for MFE estimation.

### Uridine Depletion

This constraint ensures that there is no Uridine on a third position of all the codons in an optimized sequence. This constraint is implemented on the base of DNAChisel’s CodonSpecification class.

### Dinucleotides, Codon Pair, CAI and MatchCodonUsage Optimizations

Dinucleotides and codon-pair are custom objectives which have been implemented based on usage tables taken from CoCoPUTs database^16^. These objectives account for the difference between dinucleotide or codon-pair frequencies in the host organism (*Homo sapiens* or *Mus musculus*) and the current sequence. The score is then calculated by the following formula:

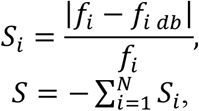

where *S*_*i*_ is the score for a given nucleotide pair or codon-pair, is the total score being the mean of all the individual scores, *f*_*i*_ is the frequency for a given pair, *f*_*i dp*_ is a corresponding frequency from the database and *N* is the total number of pairs across the sequence. The total score is used by DNA Chisel optimization algorithm as the subject for maximization.

Codon Adaptation Index (CAI) optimization is the built-in objective, which is used if a user specifies so. CAI optimization is a common optimization strategy introduced in^14^.

MatchCodonUsage constraint is a built-in constraint which minimizes the sum of discrepancies over all possible codon frequencies in a given sequence and in the target organism. This objective is set as the default one in case no other codon optimization strategies are chosen.

All codon optimization objectives are considered mutually exclusive, so it is not possible to use any combination of these in our tool.

### MFE_optimization

We are targeting to maximize the minimal free energy (MFE) at the specified region of the sequence starting from the 5’-end of the mRNA molecule. We call this region an entropy window. Maximization of MFE in this region enforces a more open structure with fewer base-pairs formed, which makes it more accessible to ribosomes. The aim is to have the MFE of the 5’-end as close to 0 as possible (it is usually negative). The user can choose between two algorithms for MFE-estimation. The first one is the RNAfold algorithm^19^, based on dynamic programming which thoroughly explores all possible secondary structures. This process can take up to several seconds depending on the size of the sequence of interest and might not be the best option when multiple runs are required (which is exactly the case of mRNAid). However, the main benefit of the long computational time is the high accuracy of estimations. RNAfold package is also used to provide the calculated secondary structures to the frontend for subsequent visualization.

The alternative option is to use a faster MFE estimation algorithm, which is based on correlated stem-loop prediction approach proposed in^20^. In this approach all possible single stem-loop conformations are considered, and their interaction energies are averaged. This algorithm has quadratic complexity O(n^2), where ‘n’ is the number of nucleotides in sequence, compared with cubic complexity O(n^3) of the RNAfold algorithm. The simplified algorithm is used during the optimization, when mutation space is explored to estimate the score of the mutated sequence. However, when presenting the final value of the best sequence after the optimization is done, its MFE value is estimated with RNAfold algorithm.

### Scoring function

Once the optimization is done and a list of optimized sequences is generated, they are ranked in order. To do so, we developed a scoring function which evaluates sequences for different criteria and assign a score for each, so that the final score looks as follows:

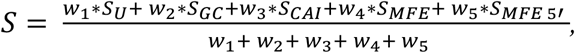

where *w*_*i*_ – are individual weights of each score (*w*_1_ = 10,*w*_2_ = 8, *w*_3_ = 7, *w*_4_ = 3, *w*_5_ = 1), *S*_*U*_ – uridine depletion score, *S*_*GC*_ – GC content score, *S*_*CAI*_ – codon adaptation index score, *S*_*MFE*_ – total MFE score, *S*_*MFE 5′*_ - MFE 5’-end score.

### Uridine depletion score

Uridine depletion is checked by counting each uridine at third position in a codon and normalizing to codon number. Maximum and minimum values are 1 and 0 (all/no codons have uridine at third position). When uridine depletion is not specified by the user, this is not included in the final scoring function (by setting the weight to 0).

### GC Score

GC content is calculated for the whole sequence and checked to be within user defined range (GC_min and GC_max). The score is calculated as a growing linear function of GC content value to favor sequences with larger GC value:

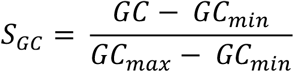

The score is bounded in the range 0 to 1. As GC content has an influence on properties and expression rates of mRNA, we optimize the sequence to fit the GC content in a specified window.

### Codon Adaptation Index score

This score is calculated as being equivalent to the value of CAI itself. This value is bounded in the range 0 to 1.

### Total MFE Score

It is preferable to have sequences with lower value of MFE. To sort the sequences according to this requirement we use the following score:

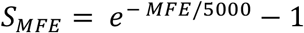

It tends to zero when MFE goes to zero, and does not exceed 1 for MFE values around <=3500 bp.

### 5’-MFE score (mfe_5_score)

MFE of the 5’-end is calculated using RNAfold. MFE has a theoretical maximum of 0, but in practice does not reach that value. The score is calculated as a decreasing exponential function of the 5’-MFE:

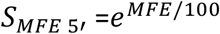

In this case s -> 0 when MFE -> minus infinity and s -> 1 when MFE -> 0. The score is now bounded in the range 0 to 1, with 0 being the worst case and 1 being the best case. The aim is to have the 5’-MFE as close to 0 as possible. This means that there are few bonds among nucleotides of the 5’-end, therefore the structure is open and more accessible to ribosomes. This is believed to improve expression.

### Built-in specifications

We also incorporated to mRNAid a set of built-in DNA Chisel specifications. Constraints include AvoidPattern that avoids a specified sequence of nucleotides, EnforceGCContent that controls the GC content across the sequence, EnforceTranslation that makes sure that the translation to a given polypeptide is preserved. Objectives include MatchTargetCodonUsage that ensures that the frequencies of codons are close to the ones in the host organism (in case we do not use other frequency optimizations, like MatchTargetPairUsage or MatchTargetCodonUsage) and EnforceGCContent that controls the GC content across the specified sliding window.

### Comparison to COVID-19 vaccines

The sequence for the spike surface glycoprotein gene was retrieved from NCBI (NC_045512.2) and then optimized by mRNAid 10 times under each software setting (Supplementary Table S1). The resultant optimized sequences were systematically compared to the relevant segment within the Pfizer/BioNtech and Moderna assembled vaccine sequences^27^. Pfizer/BioNtech and Moderna assembled vaccine sequences were downloaded from: https://github.com/NAalytics/Assemblies-of-putative-SARS-CoV2-spike-encoding-mRNA-sequences-for-vaccines-BNT-162b2-and-mRNA-1273. Following CAI optimization, the mean Levenstein distance between a given mRNAid-optimized sequence and Moderna or Pfizer/BioNtech assembled vaccine sequences was 131 and 350, respectively, reflecting approximately 3.5% and 9.1% sequence variation, when 3819 nucleotides of the spike CDS are considered (excluding the stop codon).

### Experimental Validation

#### *in vitro* transcription

mRNAs with ARCA or CleanCap® with or without Uridine modification were *in vitro* transcribed using the mMESSAGE mMACHINE® T7 Ultra transcription kit (Ambion, AMB13455). Linearized plasmid DNA containing the target gene downstream of a T7 RNA polymerase promoter was used as the template and synthesis reactions were performed according to the manufacturer’s protocol. For mRNAs with CleanCap®, T7 2X NTP/ARCA was substituted with 8 mM CleanCap® Reagent AG (TriLink Biotechnologies, N-7113) and 10 mM of each NTP. Modified uridines used included pseudouridine-5’-triphosphate (TriLink Biotechnologies, N-1019), N1-methyl-pseudouridine-5’-triphosphate (TriLink Biotechnologies, N-1081) or 5-methoxyuridine-5’-triphosphate ((TriLink Biotechnologies, N-1093). mRNAs were subsequently purified by the MegaClear Transcription Clean-up kit (Ambion, AM1908) and quantified on the NanoDrop spectrophotometer.

#### Cell culture

All cell lines were obtained from American Type Culture Collection (ATCC) and grown at 37°C, 5% CO_2_. Hepa1-6 (CRL-1830) and RAW264.7 (TIB-71) were maintained in DMEM with high glucose and GlutaMAX™ supplement (Gibco), and 10% fetal bovine serum (HyClone). The same was used for MIA PaCa-2 (CRL-1420) but with additional 2.5% horse serum (Gibco). HepG2 (HB-8065) and BJ fibroblasts (CRL-2522) were cultured in MEM with GlutaMAX™ supplement (Gibco) and 10% fetal bovine serum (HyClone). AML12 (CRL-2254) were maintained in DMEM:F12 (Gibco), 10% fetal bovine serum (HyClone), 10 µg/ml insulin, 5.5 µg/ml transferrin, 5 ng/ml selenium (Sigma I-1884) and 40 ng/ml dexamethasone (Sigma D-8893).

#### Realtime NanoLuc expression assay

Lipofectamine™ MessengerMAX™ (Life Technologies) was diluted in opti-MEM to the desired working concentration and dispensed onto 384-well white assay plates (Greiner 781080). A source plate (Labcyte LP-0200) containing serial dilutions of the mRNAs was prepared using the Bravo liquid handler (Agilent) and a 10-point 2-fold dose-titration of each mRNA was dispensed onto the assay plate using Echo (Labcyte). After a 10 min incubation, 4,000 MIA PaCa-2 or 2,000 Hepa1-6 or 3,000 AML12 or 4,000 HepG2 or 3,000 RAW264.7 or 6,000 BJ cells were added per well followed by 20 µM Endurazine (Promega), an extended time-released live cell substrate. Luminescence was measured continuously at 1-hour intervals on the Tecan Spark 10M set to 37ºC, 5% CO_2_.

#### Western blot analysis

0.08 million MIA PaCa-2 cells were seeded per well in a 24-well poly-D-lysine coated cell culture plate (Greiner) and allowed to attach overnight before mRNA transfection with Lipofectamine™ MessengerMAX™ (Life Technologies) according to the manufacturer’s protocol. After 24 hours incubation, 100 μl of Bolt™ LDS sample buffer supplemented with Bolt™ sample reducing agent was added per well of a 24-well plate. The wells were scrapped using wide orifice tips and the lysate was transferred into PCR-strip tubes and sonicated for 10 × 10 seconds in a chilled water bath sonicator (QSonica). 15 μl of protein extract was separated on 4-12% Bis-Tris plus gels, transferred onto nitrocellulose membranes using the Trans-Blot® Turbo™ semi-dry system (Bio-rad), and blocked for 1 hour at room temperature with Intercept™ (TBS) blocking buffer (Li-Cor). Blots were probed with the appropriate primary antibodies overnight at 4°C in blocking buffer supplemented with 0.1% Tween-20, followed by the secondary antibodies IRDye® 680RD donkey anti-mouse IgG or IRDye® 800CW donkey anti-rabbit IgG (Li-Cor) for 1 hour at room temperature. Fluorescent signals were imaged and quantified using Odyssey® CLx. Primary antibodies used were: NanoLuc (Promega, N7000) and GAPDH (Cell Signaling Technology, #5174)

#### IFN-β detection in BJ fibroblasts

BJ fibroblasts were seeded in 96-well poly-D-lysine coated cell culture plates (Greiner) at 20,000 cells per well and transfected the next day with 50 ng per well of the respective mRNA using Lipofectamine™ MessengerMAX™ (Life Technologies). The supernatant was harvested 48 hours post-transfection and IFN-β levels were determined using the Bio-Plex Pro Human Inflammation Panel 1 (BioRad) as per manufacturer’s protocol. Data was acquired on the Bio-Plex Pro 200 system (BioRad).

## Data availability

All code for this publication is available in the following GitHub repository: https://github.com/Merck/mRNAid and as a web application at https://mrnaid.dichlab.org.

## Author Contributions

SL, PG, BH, AP and DB conceived the study. DB planned and supervised the study. NV designed and orchestrated the implementation of mRNAid. KB and XW contributed to the development of the backend, SP to the development of the frontend, MS to the architecture and open-source, and PM to code-review and scoring function. AG acted as the product owner and managed the backlog. AM helped with scientific research throughout particularly in the context of MFE. SL and PG scientifically led and conducted the experimental work with the help of CY and JW. DB wrote the manuscript, SL, PG, JW, AM and NV also contributed to main text and method sections. All authors read and approved the final version of the manuscript.

## Acknowledgments

We thank Jens Christensen, Vincent Antonucci, and Carol A. Rohl for supporting this work. We are immensely grateful to David Dzamba for his help with the initial research on transcript stability and expression that eventually was not included in the final manuscript.

