## Supplementary Figure for "mRNAid, an Open-Source Platform for Therapeutic mRNA Design and Optimization Strategies"

**Supplementary Material**

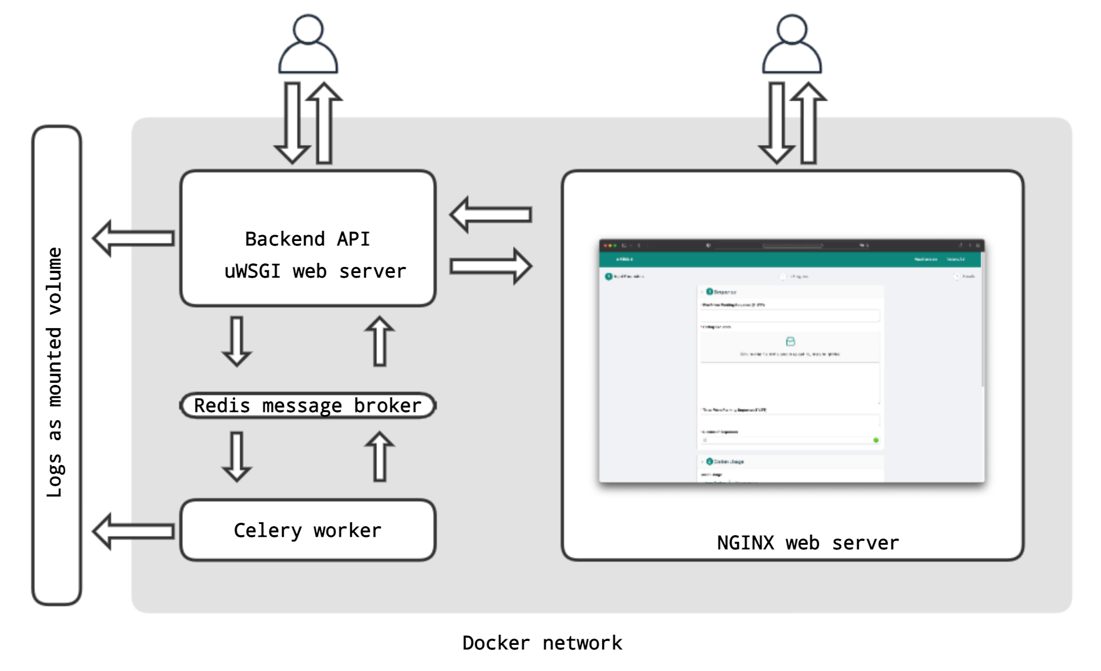

**Supplementary Figure S1**. Tool architecture. The mRNAid application consists of several parts: an Nginx web-server, a backend API, a Redis message broker and a Celery worker. All services are containerized as docker containers. Log file is mounted to the Celery worker and backend API containers.

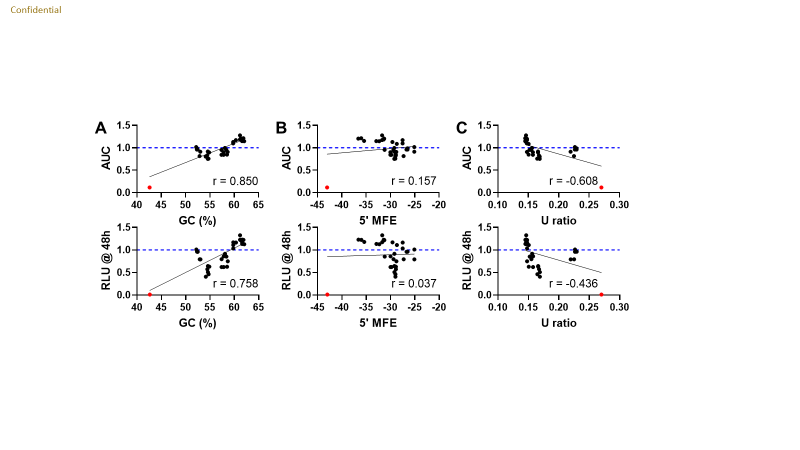

**Supplementary Figure S2.** Correlation plots for AUC (top) and RLU @ 48 hr (bottom) versus (A) GC (%), (B) MFE (kcal/mol) at the 5’-end (5’ MFE), and (C) Uridine ratio (U ratio). AUC and RLU @ 48 hr were determined for the 6.25 ng mRNA dose (Figure 1A and 1B) and represented as fold change over the Promega control (blue dashed line). The red dot denotes the de-optimized Rare input. Pearson ‘r’ is indicated as determined by GraphPad Prism.
